# Sexual identity of enterocytes regulates rapamycin-mediated intestinal homeostasis and lifespan extension

**DOI:** 10.1101/2021.10.22.465415

**Authors:** Jennifer C Regan, Yu-Xuan Lu, Enric Ureña, Ralf Meilenbrock, James H Catterson, Disna Kißler, Linda Partridge

## Abstract

Pharmacological attenuation of mTOR by rapamycin and other compounds presents a promising route for delay of ageing-related pathologies, including intestinal cancers. Here, we show that rapamycin treatment in *Drosophila* extends lifespan in females but not in males. Female-specific, age-related gut pathology and impaired intestinal barrier function are both markedly slowed by rapamycin treatment, mediated by increased autophagy. Upon rapamycin treatment, female intestinal enterocytes increase autophagy, via the H3/H4 histone-Bchs axis, while male enterocytes show high basal levels of autophagy that do not increase further upon rapamycin treatment. Sexual identity of enterocytes alone, determined by the expression of *transformer^Female^*, dictates sexually dimorphic cell size, H3/H4-*Bchs* expression, basal rates of autophagy, fecundity, intestinal homeostasis and extension of lifespan in response to rapamycin. This study highlights that tissue sex determines regulation of metabolic processes by mTOR and the efficacy of mTOR-targeted, anti-ageing drug treatments.

## Main

Sex differences in lifespan are almost as prevalent as sex itself ^1,2^. Women are the longer-lived sex in humans, in some countries by an average of >10 years, and yet bear a greater burden of age-related morbidities than do men ^3^. Many aspects of human physiology that affect homeostasis over the life course show profound sex differences, including metabolism ^4^, responses to stress ^5^, immune responses and auto-inflammation ^6–8^ and the rate of decline of circulating sex steroid hormones (menopause and andropause) ^9^. These physiological differences lead to different risks of developing age-related diseases, including heart disease, cancer, and neurodegeneration ^10,11^. Sex differences can also determine responses to pharmacological treatments ^12^; potentially both acutely, by regulating physiology and metabolism, and chronically, by influencing the type and progression of tissue pathology. Understanding how sex influences both development of age-related disease and responses to treatment will be key to move forward with the development of geroprotective therapeutics.

Greater longevity in females than in males is prevalent across taxa ^1,2,13^. Evolutionary drivers for sex differences in longevity include mating systems, physical and behavioural dimorphisms and consequent differences in extrinsic mortality, sex determination by heterogametism, and mitochondrial selection ^1,2,13,14^. Studies in laboratory model systems can help uncover the mechanisms leading to sexual dimorphism in longevity. Lifespan is a malleable trait, and genetic, environmental and pharmacological interventions can ameliorate the effects of ageing. These interventions often target highly conserved, nutrient-sensing signalling pathways, and their effects are frequently sex-specific ^12,15^. We have previously shown that dietary restriction (DR) extends lifespan more in female than in male *Drosophila melanogaster*, at least in part by targeting a dimorphic decline in gut physiology, which is much more evident in females ^16^. DR influences nutrient sensing pathways such as IIS/mTOR, and targeting these pathways directly offers a more translational route for anti-ageing therapy than do chronic dietary regimes ^17–20^.

mTOR is a highly conserved signalling hub that integrates multiple cues to regulate key cellular functions, including cell growth, division, apoptosis, and autophagy. The mTOR complex 1 (mTORC1) is activated by both nutrients and growth factors such as EGF and IIS, via PI3K and Akt, such that it responds to both organismal and intracellular energy status ^21^. Attenuation of mTORC1 activity genetically by a null mutation in the mTORC1 substrate Ribosomal protein *S6 kinase beta-1* (*S6K1*) gene increases lifespan in female, but not male, mice ^22^. Pharmacological inhibition of mTORC1 by rapamycin is currently the only pharmacological intervention that extends lifespan in all major model organisms ^17,19,23^. Rapamycin extends lifespan in mice, but the effects are also sexually dimorphic ^24^. Chronic treatment of genetically heterogenous mice, tested at 3 locations, showed moderate lifespan extensions ^24,25^, where the magnitude of extension differed substantially between the sexes. Interestingly, a subsequent study demonstrated sexually dimorphic effects on ageing pathologies, specifically cancer incidence and type ^26^. The physiological bases for these dimorphic responses to mTOR-attenuation are not well-understood. Chronic treatment with rapamycin extends lifespan significantly more in female *Drosophila melanogaster* than in males ^27^, and attenuates development of age-related gut pathologies in *Drosophila* females ^28^. However, the effect of rapamycin on ageing pathology in *Drosophila* males is unknown.

Here, we show that treatment with rapamycin extends lifespan in female flies only. Intestinal ageing in females is attenuated by rapamycin treatment, through up-regulation of autophagy in enterocytes. There are strong dimorphisms in baseline metabolic regulation of intestinal cells, whereby male enterocytes appear to represent an intrinsic, minimal limit for cell size and an upper limit for autophagy, neither of which are pushed further by rapamycin treatment. By manipulating genetic determination of tissue sex, we show that sexual identity of enterocytes determines physiological responses to mTOR attenuation, including homeostatic maintenance of gut health and function, and lifespan, through autophagy activation by the histones-Bchs axis ^29^. These data show the importance of cellular sexual identity in determining baseline metabolism, consequent rates of tissue ageing, and responses to anti-ageing interventions.

## Results

### Rapamycin treatment extends lifespan in females but not in males

We treated *w^Dah^* adult flies of both sexes with 200 μM rapamycin added to the food medium. At this dose, females, as expected ^27^, showed a significant increase in lifespan, while males did not (Fig 1a). Given that male flies eat less than females ^30,31^, and hence may ingest less of the drug, we fed females and males rapamycin at three concentrations, 50, 200 and 400 μM, in the food medium. Females showed significantly extended lifespan at all three doses of the drug (Fig S1), but males showed no increase at any dose (Fig 1b). To test whether the sexually dimorphic response of lifespan to rapamycin treatment generalised across fly genotypes, we tested the response in the *Dahomey* (*Dah*) line (from which *w^Dah^* was originally derived), and in a genetically heterogenous fly line derived by in-crossing all lines that make up the *Drosophila Genetic Resource Panel* (*DGRP-OX*) ^32^. Similar to *w^Dah^*, we observed significant lifespan extension in females, but not in males in both of these lines (Fig S2a,b). Inhibition of mTOR by rapamycin may, therefore, confer a beneficial effect in females that is absent in males. Alternatively, any beneficial physiological effect(s) in males may be counteracted by negative effects, resulting in no net change to lifespan, or males may be unable to respond to rapamycin treatment. To determine if male tissues are sensitive to inhibition of mTORC1 by rapamycin, we measured phosphorylated S6K (p-S6K) levels in dissected intestines (Fig 1c) and fat body tissue (Fig 1d) in males and females. Both sexes showed a significant reduction in p-S6K levels in both tissues in response to rapamycin, and there was no significant interaction between sex and treatment. The dimorphic response of lifespan to rapamycin was therefore likely not due to sex differences in suppression of mTORC1 signalling by the drug.

**Figure 1.**
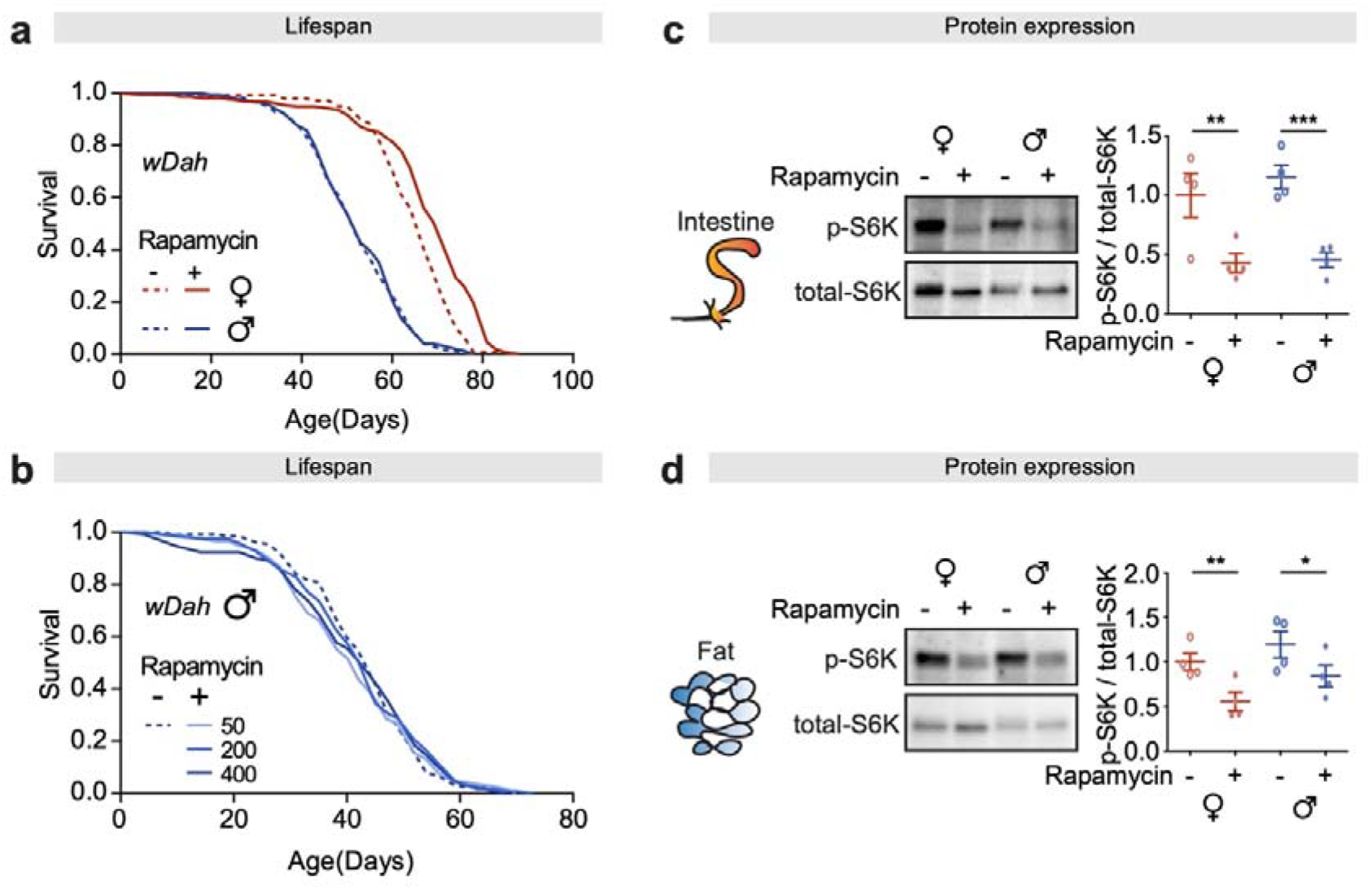
Rapamycin treatment extends lifespan in *w^Dah^* females only, but reduces phosphorylation of S6K in both sexes. **a,** Adult-onset rapamycin treatment (200 μM) extended the lifespan of *w^Dah^* females but not males (log-rank test, females p=2.1E-06, males p=0.77, n>140 flies). See also Table S1 for Cox PH analysis. **b,** Adult-onset rapamycin treatment at three concentration (50, 200 and 400 μM) did not extend the lifespan of *w^Dah^* males (log-rank test, 50 μM p=0.60, 200 μM p=0.75, 400 μM p=1, n >110 flies). See also Table S2. **c-d,** The level of phospho-S6K in the intestine and the fat body was substantially reduced by rapamycin treatment both in females and males. (n = 4 biological replicates of 10 intestines per replicate, two-way ANOVA, interaction p>0.05; post-hoc test, *p<0.05, **p<0.01, ***p<0.001).

### Age-related gut pathology is reduced in females treated with rapamycin

Dietary restriction attenuates female-specific, age-related intestinal pathologies in *Drosophila*, leading to a greater extension of lifespan in females than in males ^16^. We therefore investigated the effect of rapamycin on age-related decline in the structure and function of the gut. Small tumour formation and resulting dysplastic pathology can be quantified by assessing proportion of the intestinal epithelium which is no longer maintained as a single layer ^29,33^. In parallel with pathological changes analysed by imaging, gut barrier function can be assessed using well-described methods to detect the onset of gut leakiness ^34,35^. As previously reported ^16,28,29^, females treated with rapamycin showed a strong attenuation of dysplastic epithelial pathology (Fig 2a) and intestinal stem cell (ISC) mitoses (^36^; Fig S3a,b), in parallel with better maintenance of barrier function (Fig 2b). In contrast, male flies showed only low levels of ISC mitoses, and intestinal pathology, and these were not reduced by rapamycin treatment (Fig 2a,b; Fig S3a,b) ^37^.

**Figure 2.**
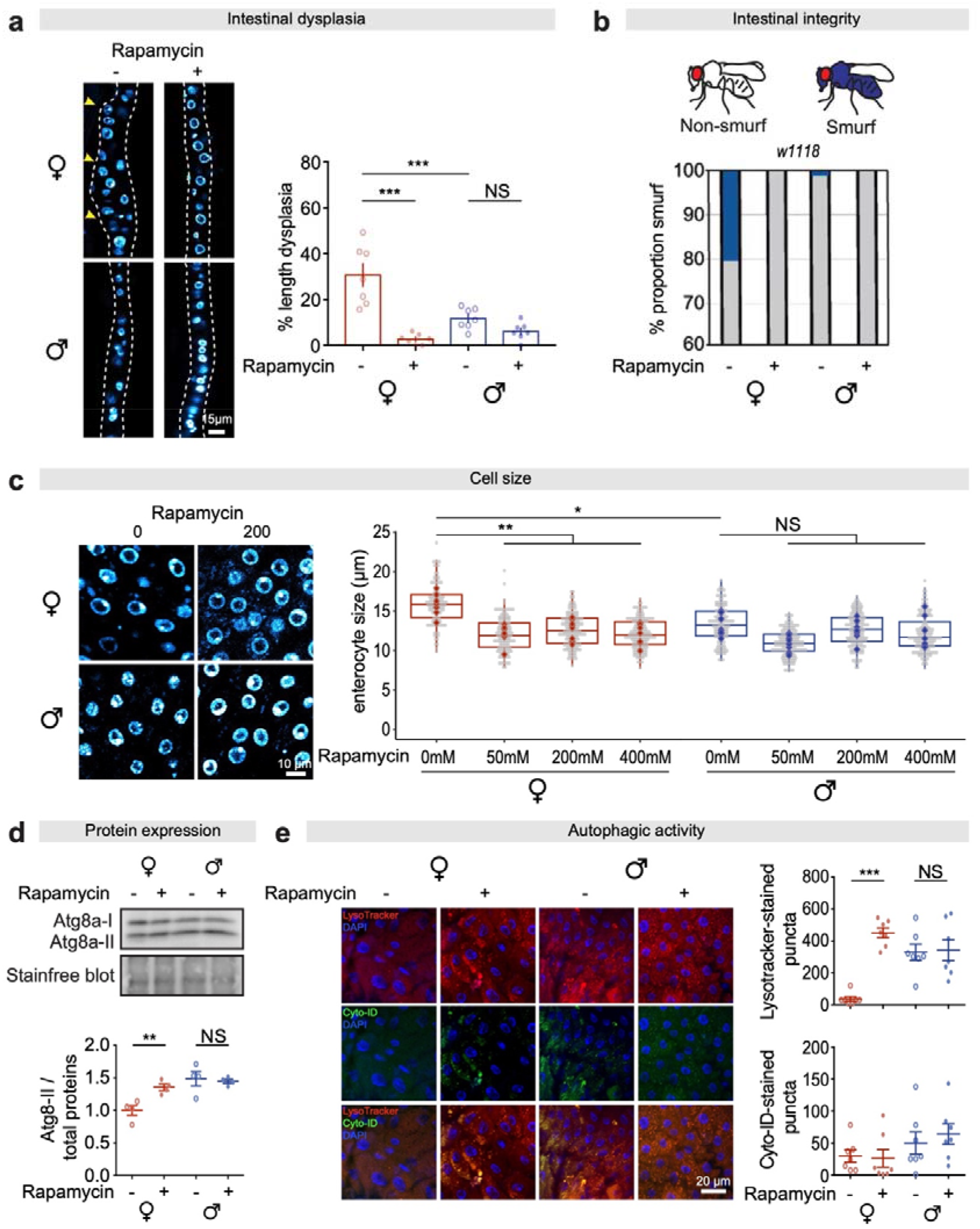
Rapamycin treatment reduces age-related gut pathology and enterocyte size, and elevates autophagy and barrier integrity in females but not in males. **a,** Females showed greater age-related dysplasia in aged guts, which was attenuated by rapamycin treatment, at 50 days of age. (n = 7 intestines, two-way ANOVA, interaction ***p<0.001; post-hoc test, ***p<0.001). **b,** Females had a higher number of flies suffering barrier function decline (Smurfs) than males, and showed increased barrier function in response to rapamycin, at 60 days of age. (n>150, Fishers exact test). **c,** Cell size of enterocytes in females was larger than in males, and declined to the same size as in males in response to rapamycin treatment (50, 200 and 400 μM) (n = 6-8 intestines, n ≥ 10 enterocytes per intestine, circles indicate individual values and diamonds represent the average value per intestine; linear mixed model, interaction p<0.01; post-hoc test, NS p>0.05, *p<0.05, **p<0.01). **d,** The expression of Atg8a-II in the gut of females was lower than in males, and rapamycin treatment increased it to the level in males (n = 4 biological replicates of 10 intestines per replicate, two-way ANOVA, interaction p<0.01; post-hoc test, NS p>0.05, **p<0.01). **e,** The number of Lysotracker-stained puncta in the gut of females was lower than in males, and rapamycin increased it to the level seen in males. Neither sex nor rapamycin had an effect on the number of Cyto-ID-stained puncta in the intestine (n = 7 intestines per condition; n = 2-3 pictures per intestine, data points represent the average value per intestine; linear mixed model, interaction Lysotracker-stained puncta, p<0.001, Cyto-ID-stained puncta, p>0.05; post-hoc test, NS p>0.05, ***p<0.001).

### The size and composition of the microbiome is sex- and age-dependent but does not change significantly upon treatment with rapamycin

Age-related shifts in the luminal microbial community can drive epithelial pathology in female *Drosophila*, due to the expansion of pathogenic bacterial species at the expense of commensals ^38^. Attenuation of the mTOR pathway by rapamycin influences composition of the microbiome in mammals ^26^. However, recent data demonstrated that chronic rapamycin treatment did not affect the microbiome in *Drosophila* females, at least under certain laboratory and diet conditions ^39^. To investigate a role for the bacterial microbiome in mediating sex differences in the responses to rapamycin under our laboratory conditions, we deep-sequenced the gut microbiome in young- and middle-aged flies of both sexes treated chronically with rapamycin. We found significant sex dimorphisms in load (Fig S4a) and composition (Fig S4b) of the microbiota, and these interacted with age. The load in old male flies increased by an order of magnitude compared with young male flies (Fig S4a), and this increase was confirmed by quantifying *Acetobacter pomorum* transcripts relative to a *Drosophila* standard. No comparable increase was seen in females, either by assessing overall load, or load of *A. pomorum*. Rapamycin treatment did not significantly affect either load (Fig S4a) or composition (Fig S4b) in either sex, suggesting that the sexually dimorphic effects of rapamycin treatment were not achieved through remodelling of the microbiome.

### Intestinal cell size is reduced in females but not in males following rapamycin treatment

TOR plays a central role in regulating antagonistic anabolic and catabolic processes, and inhibition by rapamycin concomitantly decreases cell size and up-regulates autophagy ^40,41^. We fed rapamycin at doses between 50 μM and 400 μM in the food medium and measured cell size after two weeks of treatment (Fig 2c). Enterocyte size in untreated males was significantly smaller than in untreated females, as expected ^16^, and was not significantly responsive to rapamycin treatment (Fig 2c). In contrast, treatment at 50mM reduced enterocyte size in females, to a size approximately 75% of that in control females and very similar to that in untreated males (Fig 2c), with no further reduction at 4x (200mM) or 8x (400mM) higher doses. Enterocyte size in females thus reached a minimum size, similar to that in untreated males, at a relatively low dose of rapamycin.

### Enterocytes in males have higher levels of basal autophagy that are not further increased by rapamycin treatment

Inhibition of mTORC1 by nutrient starvation, stress, or pharmacological inhibition increases autophagy ^21,40^. Autophagy can be measured *in vivo* in several ways, including Western blot analysis of the lipidated form of the Atg8a protein (Atg8a-II), the fly ortholog of mammalian LC3. There was a striking sex dimorphism in basal levels of autophagy, with Atg8a-II protein levels higher in dissected intestines from untreated males than females (Fig 2d). Rapamycin treatment substantially increased Atg8a-II in female intestines to levels similar to those in untreated males, while it had no significant effect on males (Fig 2d). To further confirm this result, we performed co-stainings with Lysotracker and Cyto-ID, which selectively label autophagic vacuoles, to assess the autophagic flux. An increased number of Lysotracker puncta indicates that autophagic flux is increased or blocked, while an increase in the number of Cyto-ID puncta indicates that flux is blocked ^29,42,43^. The number of Lysotracker-stained puncta in untreated female intestines was significantly lower than in males (Fig 2e), and when treated with rapamycin increased to levels that did not differ significantly from the basal level in males, while there was no increase in male intestines (Fig 2e). Neither sex nor rapamycin treatment affected the number of Cyto-ID puncta (Fig 2e), suggesting autophagic flux was not blocked. Taken together, these results demonstrate that males had higher basal levels of autophagy than did females, and that only in females was there an increase in response to rapamycin, which brought autophagy to similar levels to those seen in males.

### Suppressing autophagy in enterocytes reduces barrier function and decreases lifespan in males

To probe the role of increased basal autophagy levels in males, we genetically suppressed the process, by expressing RNA interference (RNAi) against the essential autophagy gene *Atg5* in adult enterocytes, using the Geneswitch system ^44^; *5966GS>Atg5^[RNAi]^*. In line with our previous result (Fig 2e), males showed markedly higher basal levels of intestinal autophagy than did females (Fig 3a). Knock-down of *Atg5* reduced the number of Lysotracker-stained puncta in males to similar levels as in females, while females showed no response (Fig 3a).

**Figure 3.**
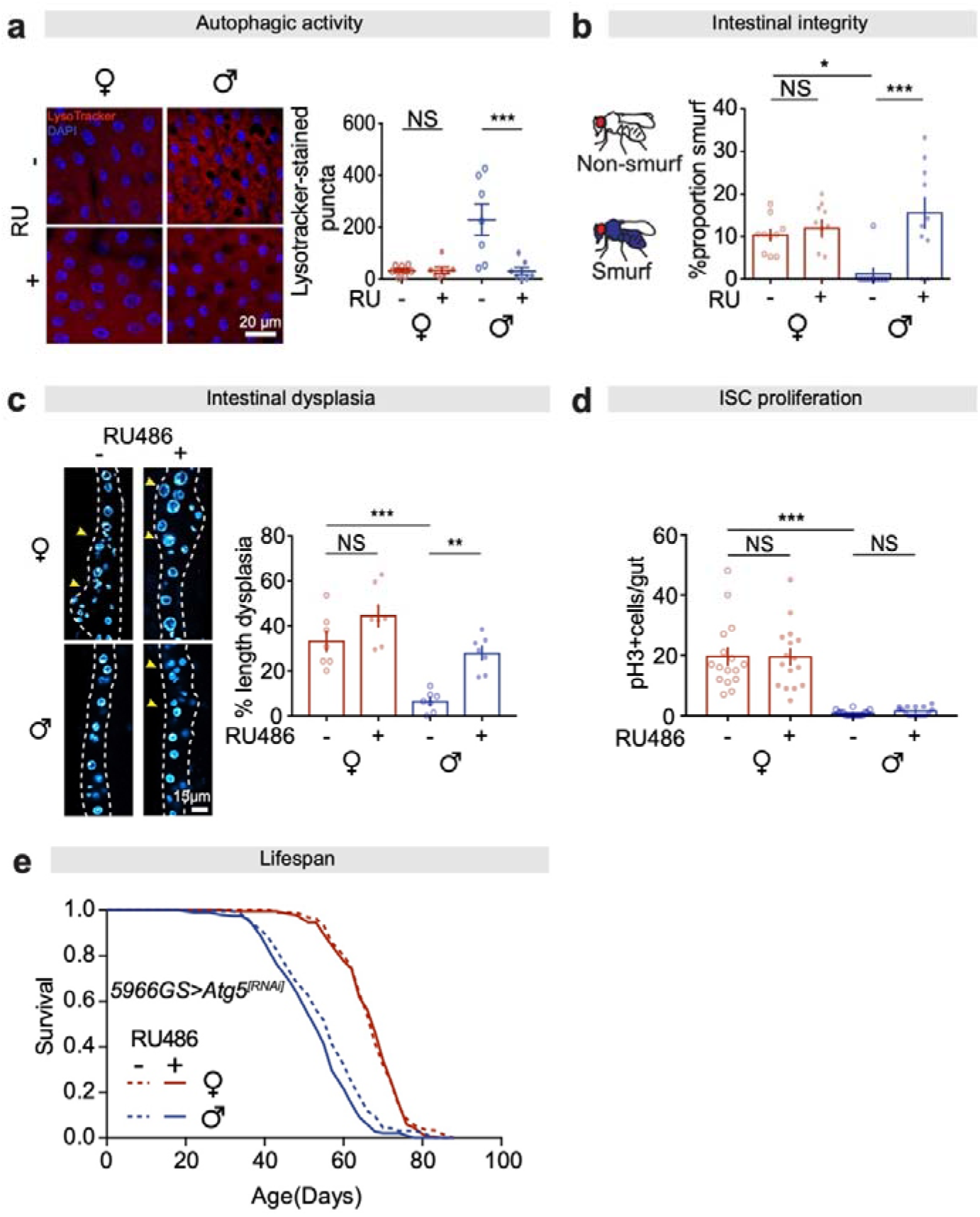
Autophagy in gut enterocytes regulates gut pathologies and lifespan. **a,** Adult-onset knock-down of *Atg5* in adult ECs did not affect the number of Lysotracker-stained puncta in the gut of females, but decreased it in the gut of males to the level in females, at 20 days of age. (n = 7 intestines per condition; n = 2-3 pictures per intestine, data points represent the average value per intestine; linear mixed model, interaction p<0.01; post-hoc test, NS p>0.05, ***p<0.001). **b,** Females had a higher number of Smurfs than males, and adult-onset knock-down of *Atg5* in adult ECs in males significantly increased the number of Smurfs, to the level in females at 60 days of age. Bar charts show n = 10 biological replicates of 8-20 flies per replicate (two-way ANOVA, interaction p<0.01; post-hoc test, NS p>0.05, *p<0.05, ***p<0.001). **c,** Adult-onset knock-down of *Atg5* in adult ECs did not affect the level of dysplasia in the gut of females, but increased it in the gut of males to the level in females, at 50 days of age. (n = 7 intestines, two-way ANOVA, interaction p>0.05; post-hoc test, NS p>0.05, **p<0.01, ***p<0.001). **d,** Adult-onset knock-down of *Atg5* in adult ECs did not change the number of pH3 + cells in either females or males, at 20 days of age. (n = 16 intestines, two-way ANOVA, interaction p>0.05; post-hoc test, NS p>0.05, ***p<0.001). **e,** Adult-onset knock-down of *Atg5* in adult ECs shortened lifespan of males but not females (log-rank test, females p=0.80, males p=4.5E-03, n≥195 flies). See also Table S4 for Cox PH analysis.

Autophagy maintains homeostasis of ageing tissues, and its manipulation can affect lifespan ^45,46^. Indeed, gut barrier function was reduced in aged male flies with suppressed autophagy, to levels similar to those seen in females (Fig 3b). In contrast, expression of *Atg5^[RNAi]^* had no effect on barrier function in female flies (Fig 3b), likely due to the lack of response of their already low levels of intestinal autophagy to knock-down of *Atg5*. Development of dysplasia was also significantly increased in aged *5966GS>Atg5^[RNAi]^* males compared to controls, but again there was no significant effect in females (Fig 3c). Interestingly, when we analysed ISC proliferation in 20-day old flies, we did not see an up-regulation of mitoses in male flies with suppressed autophagy in enterocytes (Fig 3d). This suggests that the dysplasia we observed in these flies was the cumulative effect of disrupted differentiation of ISCs or enteroblasts, arising as a non-cell autonomous effect of decreased autophagy in neighbouring enterocytes, rather than a consequence of increased proliferation. RNAi against *Atg5* in enterocytes significantly decreased lifespan in male flies, but had no effect in females (Fig 3e). These data reveal the dimorphic regulation of autophagy in enterocytes and its impact on gut pathology and lifespan: females have low basal levels autophagy which increase in response to rapamycin treatment, with a consequent reduction in gut pathology and increase in lifespan, whereas males with high basal autophagy see an increase in gut pathology and a reduction in lifespan upon its suppression.

### Cellular and molecular responses to TOR-attenuation depend on cell-autonomous sexual identity of enterocytes

In *Drosophila*, somatic cells determine sexual identity in a cell-autonomous manner, based on sex chromosome karyotype, via the sex determination pathway ^47^. Genetic manipulation of the pathway at the level of the splicing factor *transformer* allows for the generation of tissue-specific sexual chimeras ^16,48^. To test the role of cell-autonomous sexual identity in regulating sexually dimorphic phenotypes, we switched sex solely in enterocytes, of males and females, through the expression or abrogation of *transformer^Female^* (*tra^F^*).

Enterocyte size is regulated both by sex and TOR-signalling (Fig 2c). Masculinisation of female cells through enterocyte-specific expression of *tra^F[RNAi]^*, reduced cell size to that of males, and this was not reduced further by treatment with rapamycin (Fig S5b). In contrast, feminisation of male enterocytes by expression of *tra^F^* did not affect their size, and neither did treatment with rapamycin (Fig S5a). This suggests that expression of *tra^F^* is necessary, but not sufficient, for the larger cell size observed in female intestines.

Using Lysotracker-staining to assess the level of autophagy in the intestines of sexual chimeras, we observed that males expressing *tra^F^* in enterocytes (*mex1Gal4;UAS-tra^F^*) had suppressed basal autophagy levels in the intestine, and this showed a significant increase upon treatment with rapamycin (Fig 4a), similar to control females. In concordance, females expressing *tra^F[RNAi]^* in enterocytes (*mex1Gal4;UAS-tra^F[RNAi]^*) had increased autophagy compared to control females, but did not respond to treatment with rapamycin (Fig 4b), similar to control males. These data suggest that levels of autophagy in enterocytes are determined by enterocyte sex, not organismal sex.

**Figure 4.**
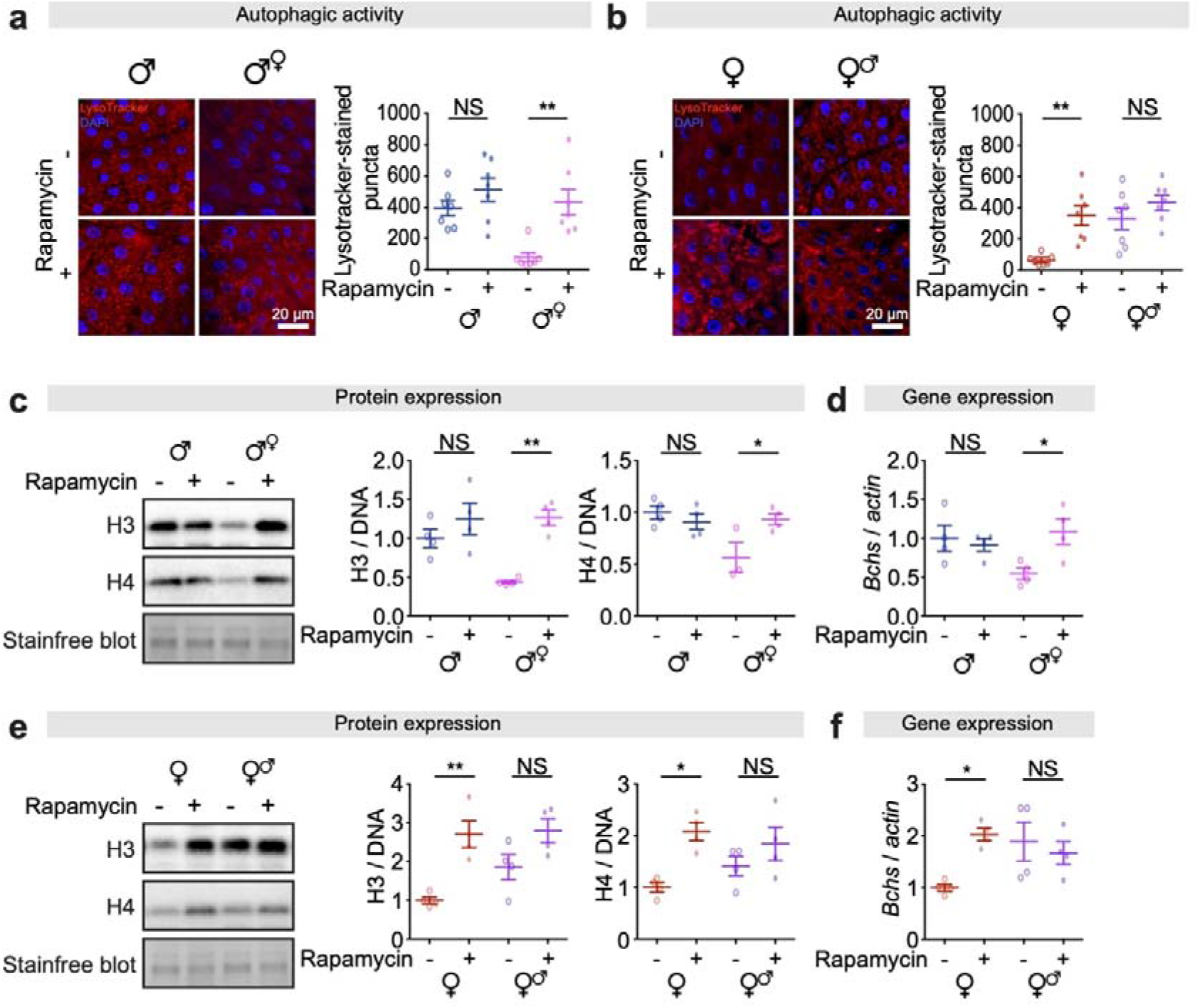
Cell-autonomous sexual identity in enterocytes dictates the levels of autophagy, histones and *Bchs* expressions in response to rapamycin treatment. **a,** Feminisation of male guts by expression of *tra^F^* in ECs reduced the number of Lysotracker-stained puncta in the gut, and it restored the response to rapamycin treatment. (n = 7 intestines per condition; n = 2-3 pictures per intestine, data points represent the average value per intestine; linear mixed model, interaction p<0.05; post-hoc test, NS p>0.05, **p<0.01). **b,** Masculinisation of female guts by knock-down of *tra^F^* in ECs increased the number of Lysotracker-stained puncta in the gut, and abolished the response to rapamycin treatment. (n = 7 intestines per condition; n = 2-3 pictures per intestine, data points represent the average value per intestine; linear mixed model, interaction p<0.05; post-hoc test, NS p>0.05, **p<0.01). **c,** Expression of histones H3 and H4 in the gut of feminised males was lower than in males, and rapamycin treatment increased it to the level in males (n = 3-4 biological replicates of 10 intestines per replicate, two-way ANOVA, H3 and H4, interaction p<0.05; post-hoc test, NS p>0.05, *p<0.05, **p<0.01). **d,** Expression of *Bchs* in the gut of feminised males was lower than in males, and rapamycin treatment increased it to the level in males (n = 4 biological replicates of 10 intestines per replicate, two-way ANOVA, H3 and H4, interaction p<0.05; post-hoc test, NS p>0.05, *p<0.05). **e,** Expression of histones H3 and H4 in the gut of masculinised females was higher than in females, and rapamycin treatment did not increase it further (n = 4 biological replicates of 10 intestines per replicate, two-way ANOVA, interaction p>0.05; post-hoc test, NS p>0.05, *p<0.05, **p<0.01).

Our recent study demonstrated that intestinal autophagy is mediated through a histones-Bchs axis, where levels of H3 and H4 histone proteins regulate the autophagy cargo adaptor *bluecheese* (*Bchs*) in enterocytes^29^. Publicly-available expression data (*FlyAtlas 2*) indicates that *Bchs* is expressed at higher levels in intestines of males than of females ^49^. We confirmed that *Bchs* transcript levels, and expression of histones H3 and H4 proteins, were higher in intestines of males compared to females. Rapamycin treatment did not increase either *Bchs* or histone expression further in males but did so in females, to levels comparable with those in males in the case of *Bchs* (Fig S6a,b). Notably, the level of H3, H4 and *Bchs* was strictly correlated with the level of autophagy in the intestines of sexual chimeras. Feminised males showed a low level of H3, H4 and *Bchs* which was increased to the same level as that of control males in response to rapamycin treatment (Fig 4c,d). Masculinised females had high basal levels of H3, H4 and *Bchs*, which were not increased further in response to rapamycin treatment (Fig 4e,f). These results suggest that the histone H3/H4-*Bchs* axis plays a key role in the sexual dimorphism of intestinal autophagy.

### Sexual identity of enterocytes influences fecundity and determines the response of intestinal homeostasis and lifespan to rapamycin

Limited cell growth and increased autophagy are correlated with better intestinal homeostasis during ageing in males compared to females (Fig 2c-e). To determine if this correlation held in individuals with sex-switched enterocytes, we measured intestinal dysplasia, barrier function, and ISC hypermitosis. In concordance with analyses of autophagy in young individuals, intestinal dysplasia and barrier function were correlated with enterocyte sex, as were responses of these pathologies to rapamycin (Fig 5a,b,d,e). ISC mitoses were affected by enterocyte sex, such that males with feminised enterocytes had higher numbers of mitoses, and females with masculinised enterocytes had fewer (Fig 5c,f). This is in line with other evidence of non-cell autonomous effects of enterocyte homeostasis on ISCs ^50^.

**Figure 5.**
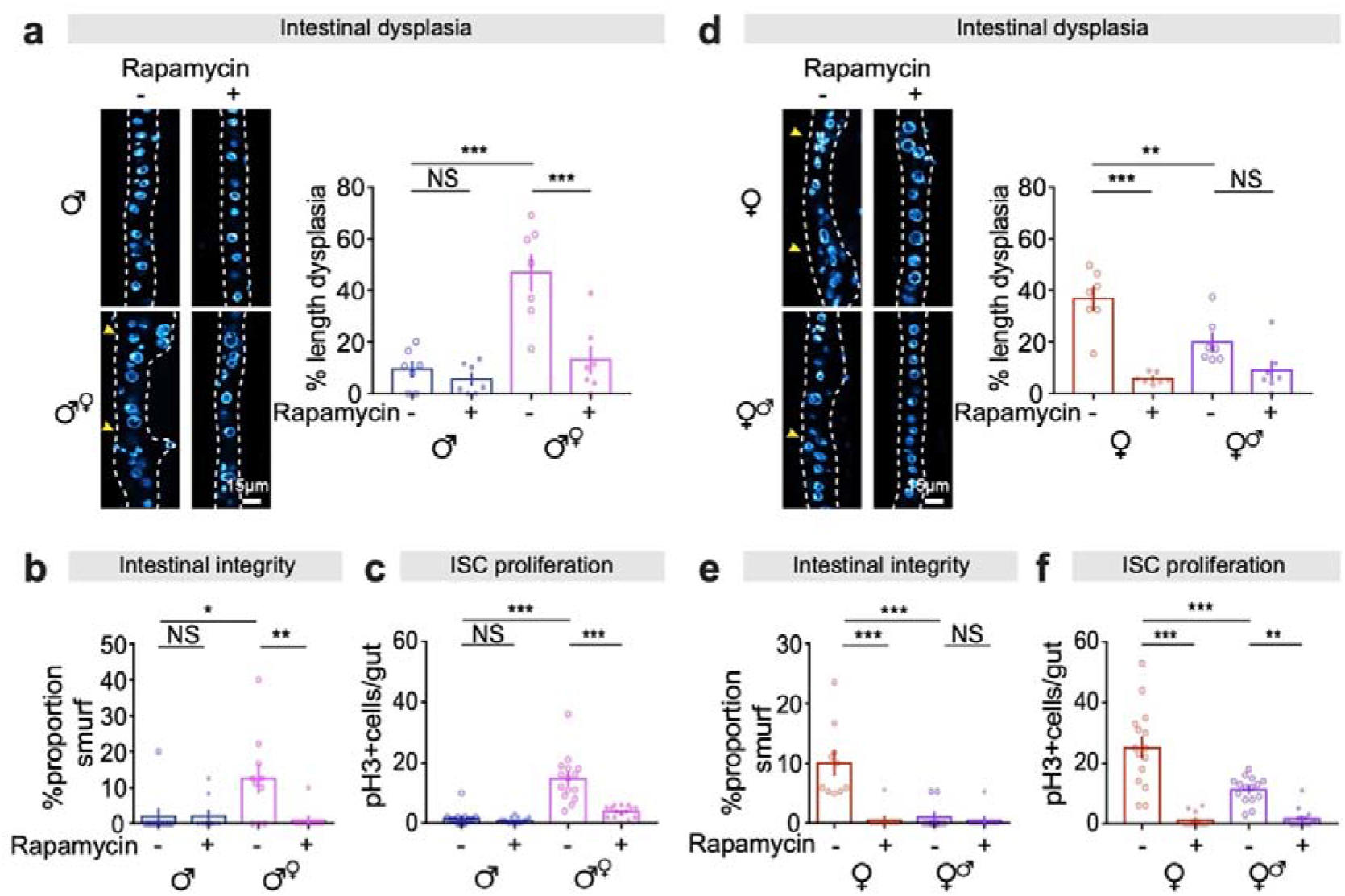
Cell-autonomous sexual identity in enterocytes mediates age-related gut pathology, barrier function and ISC mitoses in response to rapamycin treatment. **a,** Feminisation of male guts by expression of *tra^F^* in ECs increased intestinal dysplasia, which was attenuated by rapamycin treatment, at 50 days of age. (n = 7 intestines per condition; two-way ANOVA, interaction p<0.01; post-hoc test, NS p>0.05, ***p<0.001). **b,** Feminisation of male guts by expression of *tra^F^* in ECs increased the proportion of Smurfs, which was attenuated by rapamycin treatment, at 60 days of age. Bar charts show with n = 10 biological replicates of 6-12 flies per replicate (two-way ANOVA, interaction p<0.05; post-hoc test, NS p>0.05, *p<0.05, **p<0.01). **c,** Feminisation of male guts by expression of *tra^F^* in ECs increased the number of pH3 + cells, which was attenuated by rapamycin treatment, at 20 days of age. (n = 15 intestines per condition; two-way ANOVA, interaction p<0.001; post-hoc test, NS p>0.05, ***p<0.001). **d,** Masculinisation of female guts by knock-down of *tra^F^* in ECs decreased intestinal dysplasia, which was not further decreased by the combination of rapamycin treatment, at 50 days of age. (n = 7 intestines per condition; two-way ANOVA, interaction p<0.01; post-hoc test, NS>0.05, **p<0.01, ***p<0.001). **g,** Masculinisation of female guts by knock-down of *tra^F^* in ECs decreased the proportion of Smurfs, which was not further decreased by the combination of rapamycin treatment, at 60 days of age. Bar charts show with n = 10 biological replicates of 15-20 flies per replicate (two-way ANOVA, interaction p<0.001; post-hoc test, NS p>0.05, ***p<0.001). **h,** Masculinisation of female guts by knock-down of *tra^F^* in ECs decreased the number of pH3+ cells, which was further decreased by the combination of rapamycin treatment, at 20 days of age. (n = 15 intestines per condition; two-way ANOVA, interaction p<0.001; post-hoc test, **p<0.01, ***p<0.001).

Gut growth via ISC division ^48,51^, and some aspects of intestinal metabolism ^52^, have been demonstrated to impact fertility in females and males, respectively. To determine whether enterocyte sex can influence reproductive output, we measured fertility in individuals with sex-switched enterocytes. We did not detect a difference in the fertility of enterocyte-feminised males compared to that of control males (Fig 6a,b). However, enterocyte-masculinised females showed moderately, but significantly, decreased fertility, compared to that of control females (Fig 6a,c).

**Figure 6.**
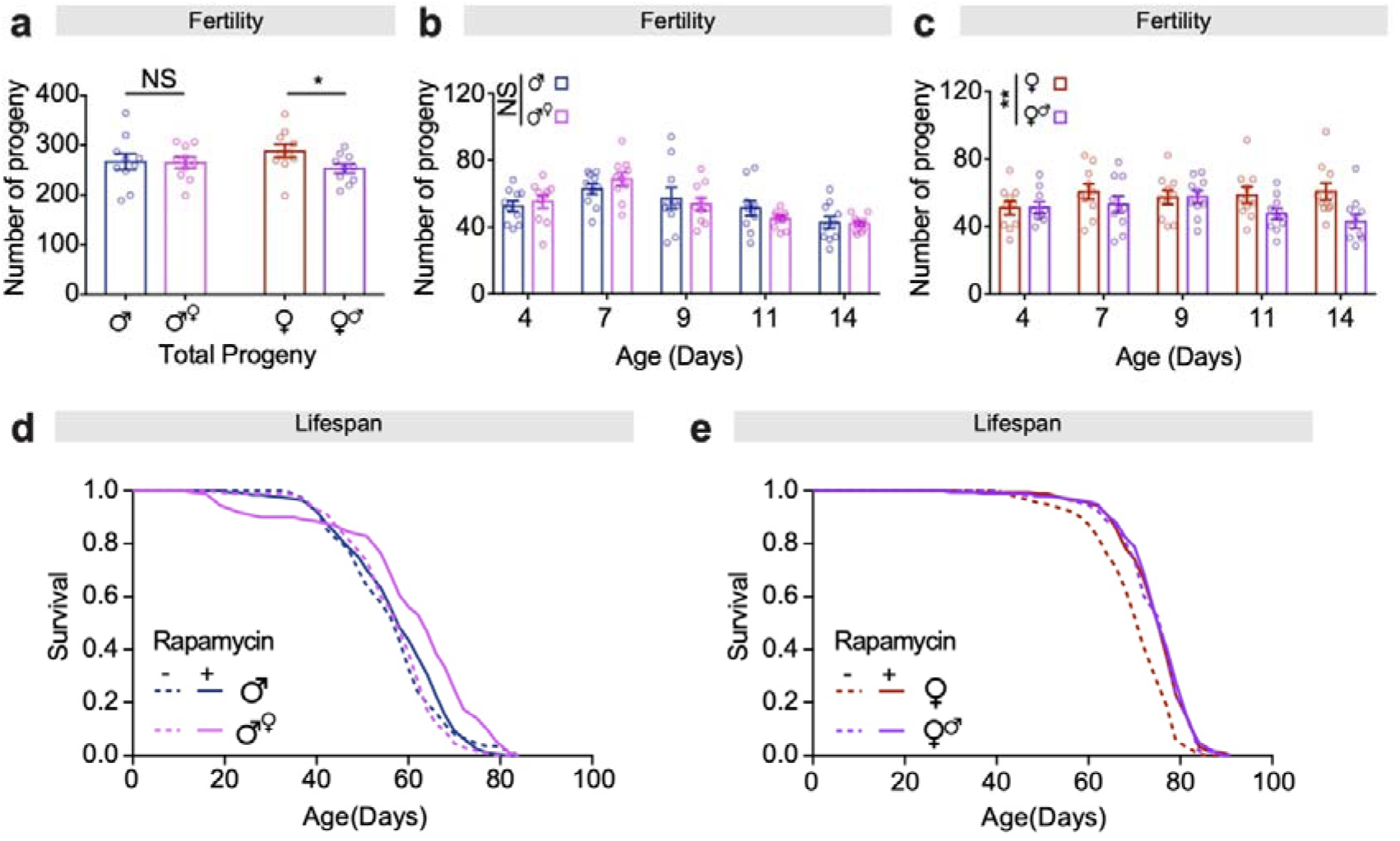
Cell-autonomous sexual identity in enterocytes influences fertility, and it mediates extension of lifespan in response to rapamycin treatment. **a-c,** Feminisation of male guts by expression of *tra^F^* in ECs did not affect the number of progeny, while masculinisation of female guts by knock-down of *tra^F^* in ECs reduced the number of progeny (n = 10 biological replicates of 3 males and 3 females per replicate, (**a**) students t test, NS p>0.05, *p<0.05; (**b**) two-way ANOVA, treatment p>0.05; (**c**) two-way ANOVA, treatment p<0.01). **d,** Feminisation of male guts by expression of *tra^F^* in ECs extended lifespan in response to rapamycin treatment (log-rank test, p= 1.55E-06, *mexG4> tra^F^* Control vs *mexG4>tra^F^* Rapamycin, n >190 flies). See also Table S5 for Cox PH analysis. **e,** Masculinisation of female guts by knock-down of *tra^F^* in ECs extended lifespan, which was not further extend by rapamycin treatment (log-rank test, p= 1.56E-09 *mexG4>+* Control vs *mexG4> tra^F_[RNAi]_^* Control, n >190 flies). See also Table S6 for Cox PH analysis.

Feminised males showed a lifespan extension upon treatment with rapamycin that was not observed in control males (Fig 6d). In contrast, masculinized females did not extend lifespan in response to rapamycin (Fig 6e). Interestingly, while the lifespan of gut-feminised males was not shorter on control food compared to that of control males (Fig 6d), the lifespan of gut-masculinized females on both rapamycin-treated and control food was comparable to that of control females treated with rapamycin (Fig 6e), suggesting an interaction between enterocyte size, enterocyte autophagy, intestinal pathology, fertility, and pharmacological mTOR-attenuation by rapamycin, which consequently mediates lifespan.

Altogether, these results suggest that the intrinsic sexual identity of enterocytes determines the effect of rapamycin on intestinal homeostasis and lifespan, where individuals with male enterocytes do not reduce intestinal pathology or extend lifespan under rapamycin treatment while flies with female enterocytes do so, regardless of organismal sex.

## Discussion

The IIS/mTOR signalling network regulates dimorphic, complex traits such as metabolism, growth, and lifespan ^22,53–55^. However, it is not well understood how dimorphisms in IIS/mTOR-regulated traits impact tissue ageing and responses to geroprotective drugs. Targeted mTORC1 inhibition by the drug rapamycin extends lifespan more in female than in male mice ^24,56^. Although there is evidence that off-target effects of rapamycin on hepatic mTORC2 signalling via *Rictor* can reduce the lifespan of male mice ^57^, dimorphic effects of rapamycin treatment on lifespan may also be regulated by other, complex interactions with specific tissues and through interaction with environmental factors such as the microbiome ^26^. Responses of lifespan to rapamycin treatment trials in mice were dose-dependent, and we do not yet know the maximum lifespan extension that can be achieved, in either sex, through chronic treatment with the drug. In one study, female mice were found to have higher circulating levels of rapamycin than did males for a given dose in the food ^24^, suggesting that sex differences in drug metabolism or bioavailability could play a role in dimorphic responses to pharmaceutical therapies ^12^. *Drosophila*, which shows a strong lifespan extension in females treated with rapamycin ^27^, offers a tractable system for understanding tissue-specific contributions to ageing dimorphisms ^16^, and dimorphic responses to anti-ageing therapeutics^19,58^.

Here, we show that treatment of *Drosophila* with rapamycin extends lifespan in females but not in males, regardless their genetic background. Rapamycin treatment increases autophagy and reduces cell size of intestinal enterocytes in females. We demonstrate a striking dimorphism in basal metabolism of enterocytes: in males, autophagy is constitutively high, cell size is smaller than in females, and both autophagy and cell size are insensitive to mTORC1-attenuation by rapamycin. This raises the possibility that intestinal autophagy is actively buffered in males, or is maintained at an upper limit by constraints on the availability of autophagy components in enterocytes. One consequence of increased intestinal autophagy in males is attenuated age-related intestinal barrier function decline, underpinning the overall slower progression of age-related intestinal pathologies in males compared to females. Intestinal barrier function maintenance, independent of ISC division, is a key determinant of lifespan in *Drosophila*. This has been demonstrated in multiple ways in females through manipulation of diet ^59^, the microbiome ^38^, and through genetic targeting of junctional components ^60^ or upstream signalling pathways ^29,61^. Males do not usually respond strongly to manipulations that attenuate functional decline of the intestine ^16,58^, including rapamycin (^27^ and this study), likely because progression of intestinal pathology is slow. Here, we show that males are also sensitive to barrier function decline, by genetically targeting autophagy components, which increased the incidence of barrier function failure, and decreased lifespan in males.

Our recent study demonstrated that a specific autophagy pathway, regulated by histones H3/H4 and requiring the cargo adaptor Bchs/WDFY3, maintains junctional integrity in enterocytes in the intestine in females during ageing^29^. Autophagy in enterocytes also lowers sensitivity to ROS induced by commensal bacteria, via suppression of p62 and Hippo pathway genes, to maintain septate junction integrity and attenuate dysplasia ^62^. Maintenance of cell junctions by increased autophagy is not restricted to epithelial tissue; for example, this occurs acutely in mammalian endothelial cells to prevent excessive diapedesis of neutrophils in inflammatory responses ^63^. Here, we demonstrate a link between enterocyte sex, the histone-Bchs axis, junctional integrity, and lifespan. We show that cell-autonomous sexual identity of enterocytes determines their histone and *Bchs* levels, and subsequently their basal level of autophagy. Autophagy is key to maintaining junctional integrity in enterocytes and, consequently, barrier function of the intestine. Thus, the sex-determined metabolic state of enterocytes, including basal autophagy and cell size, dictates how they respond to rapamycin treatment; at the cellular level, at the level of organ physiology, and at the level of whole organism homeostasis during ageing to influence lifespan ^55,64^.

Why do males and females take such different approaches to intestinal homeostasis? Females pay a cost during ageing for maintaining their intestine in an anabolic state, with lower autophagy, higher cell growth, and higher rates of stem cell division (this study, ^16,48^) leading to pathology and dysplasia at older ages ^16^. Selection acts weakly on age-related traits and strongly on those promoting fitness in youth ^65^, and females require hormone-regulated intestinal cell growth and organ size plasticity to maintain egg production at younger ages ^51,66^. Here, we show that metabolic responses of the intestine to mTOR-attenuation, including autophagy and cell growth, are regulated by *tra* cell-autonomously. Sensitivity to nutrients, particularly protein levels, in the diet is important for females to maintain and regulate egg production ^67^; we show that female enterocytes have a cell-autonomous sensitivity to changes in mTOR-signalling. This may be an adaptive mechanism to maintain reproductive output in the face of fluctuating nutrient availability ^68^, where females can take advantage of higher protein by resizing enterocytes ^69^, in addition to post-mating organ growth achieved through stem cell division ^48,51^. We show that females with masculinised enterocytes, which have a smaller cell size and higher autophagy, have reduced fertility. This is similar to the reduction in fertility demonstrated when ISCs are masculinised in female guts ^48^, suggesting that sex-determination signalling regulates organ size plasticity via both cell growth and cell division. Although fertility was reduced, enterocyte-masculinised females had healthier guts over ageing and a longer lifespan, supporting the idea that in females, early life reproduction trades-off with intestinal homeostasis at older ages ^66^.

Interestingly, males with feminised enterocytes did not show an increase in enterocyte cell size, suggesting that *tra^F^* is necessary, but not sufficient, to induce enterocyte growth, contrary to the effect seen on whole body size when *tra^F^* is expressed throughout the developing larva ^70^. Females produce larger enterocytes when flies are fed with a high protein diet, or through genetically activating mTOR or blocking autophagy by manipulation of mTOR-autophagy cascade core components in a cell-autonomous manner ^69^. However, we find that manipulating enterocyte sex, and consequently autophagy levels, does not lead to larger cells in males. Together, these data suggest that feminising enterocytes by overexpression of *tra^F^* in male guts does not simply recapitulate autophagy reduction by enterocyte-specific knock-down of *Atg5*. One possibility is that feminised enterocytes maintain better nutrient absorption during ageing, a known determining factor of lifespan ^71,72^, counteracting the effect of increased pathology and leading to comparable lifespan to males on control food.

Male fertility was unaffected by feminisation of enterocytes. Male fitness may rely more heavily on nutrients other than protein, particularly carbohydrates, where non-autonomous regulation of sugar metabolism in the male gut by the testis has been shown to be essential for sperm production ^52^. The sexes, therefore, rely on distinct metabolic programmes to maintain fitness. Cellular growth and size plasticity of the gut may not increase fitness in males, and as a result, they may maintain their intestines at a low catabolic limit that cannot be pushed further by lowered mTOR. Sexually antagonistic traits can be resolved by sex-specific regulation ^73^. Direct regulation of cell growth and autophagy (this study) and stem cell activity ^48^ by sex determination genes may allow males and females to diverge in their energetic investment in the gut, and this may interact with fertility and pathophysiology, which can eventually determine lifespan.

Importantly, the higher basal levels of autophagy in males appears to be conserved in rodents, since male mice have higher basal levels of autophagy than do females. This is seen in multiple tissues, including transcription of autophagy-related genes in spinal cord and muscle tissue ^74^, and autophagy proteins in the heart and liver ^75^, of male mice. These sex differences are present from early development and into adulthood, and are speculated to contribute to the greater female vulnerability to age-related disorders such as Alzheimer’s disease ^76^. Sex differences in baseline metabolism may profoundly influence responses to a broad range of treatments to such age-related disorders, particularly those that target nutrient-sensing pathways.

Understanding sex differential responses to geroprotective interventions gives an understanding of the mechanistic underpinnings of sex differences in the intrinsic rate of ageing in specific tissues ^14,77^, including sex-specific trade-offs. When we treat age-related disease, we are not treating individuals with equal case histories, but individuals impacted by a lifetime of differences, including those regulated by sex. Sex will be a fundamental distinction made in precision medicine, and understanding conserved mechanisms regulating dimorphism and determining responses to therapeutics will allow for the development of sex-optimised treatments.

## Acknowledgements

We thank Paula Juricic Dzankic and Jenny Fröhlich for help in preparing tissues and experiments. We thank Paulina Mika, Mary-Kate Corbally and Rebecca Belmonte for their help in maintaining lifespan experiments. We thank Oliver Hahn for his help with microbiome data analysis and the Max Planck Genome Center Cologne for performing next-generation sequencing. We thank the FACS and Imaging Core Facility at the Max Planck Institute for Biology of Ageing for their help with microscopy data. We thank Adam Dobson and David Duneau for critical reading of the manuscript and colleagues at University of Edinburgh, UCL IHA, and MPI-Age for their feedback on the study. The Bloomington Drosophila Stock Center (NIH P40OD018537) and Vienna Drosophila Resource Center (VDRC) are acknowledged for fly lines. This project has received funding from the European Research Council (ERC) under the European Union’s Horizon 2020 research and innovation programme no. 741989 and the Max-Planck-Gesellschaft. Jennifer C Regan was supported by a Wellcome Trust Seed Award (210183/Z/18/Z), and start-up funding from The University of Edinburgh. Yu-Xuan Lu was supported by an EMBO Long-Term Fellowship (ALTF 419-2014).

## Author Contributions

J.C.R., Y.X.L. and L.P. conceived the study and designed the experiments, J.C.R., Y.X.L., E.U., R.M., J.H.C. and D.K. conducted the experiments, J.C.R., Y.X.L., E.U. and R.M. analysed the data, J.C.R. and Y.X.L. wrote the original draft of paper, J.C.R., Y.X.L. and L.P. reviewed and edited the paper. Both J.C.R and Y.X.L, contributed equally and have the right to list their name first in their CV. All authors contributed to the article and approved the submitted version.

## Competing Interests

The authors declare no competing interests.

## Method Details

### Fly stocks and husbandry

All transgenic lines were backcrossed for at least six generations into the outbred line, *w^Dah^* maintained in population cages (unless specified otherwise in figure legends). Stocks were maintained and experiments conducted at 25°C on a 12 hr: 12 hr light/dark cycle at 60% humidity, on food containing 10 % (w/v) brewer’s yeast, 5% (w/v) sucrose, and 1.5% (w/v) agar unless otherwise noted. The following stocks were used in this study: *UAS-Atg5^[RNAi]^*^78,79^, *UAS-tra^F^* (Bloomington #4590), *UAS-tra^F[RNAi]^* (Bloomington #44109), *mex1Gal4* (Bloomington #91369), *5966GS* ^80^, *Dah* ^81^, *DGRP-OX*^32^.

### Lifespan assay

Files were reared at standard density before being used for lifespan experiments. Crosses were set up in cages with grape juice agar plate. The embryos were collected in PBS and squirted into bottles at 20 μl per bottle to achieve standard density. The flies were collected over a 24 h period and allowed 48 h to mate after eclosing as adults. Flies were subsequently lightly anaesthetized with CO_2_, the adults were sorted into the vials at a density of 20/vial. Rapamycin (LC laboratories) and/or RU486 (Sigma) dissolved in ethanol was added to food. For control food ethanol alone was added.

### Fertility assay

All fertility assays were performed on vials housing 3 virgin females and 3 virgin males that were all 2 days old. All assays were performed on 10 replicates per group. Flies were transferred to new vials every 2–3 days and flies were discarded after the fifth’ ‘flip’. In order to assess overall fertility, we counted emergence of pupal progeny, as previously described ^82^.

### Gut barrier assay (“Smurf” assay)

Flies were aged on normal SYA food and then switched to SYA food containing 2.5% (w/v) Brilliant Blue FCF (Sigma). Flies were examined after 48 h, as previously described ^16,34^.

### Immunoblotting

Fly tissues were homogenized in 80μl 1x RIPA Lysis and Extraction Buffer (Thermofisher) containing PhosSTOP (Roche) and cOmplete, Mini, EDTA-free Protease Inhibitor Cocktail (Roche). Extracts were then cleared by centrifugation, protein content determined by using Pierce™ BCA Protein Assay (Thermofisher) and DNA content determined by using Qubit dsDNA HS Assay (Invitrogen). Approximately 8μg of protein extract was loaded per lane on polyacrylamide gel (4-20% Criterion, BioRad). Proteins were separated and transferred to PVDF membrane. Following antibodies were used: Atg8a (Péter Nagy’s lab, 1:5000), Phospho-Drosophila p70 S6 Kinase (Thr398) (Cell Signaling #9209, 1:1000), total S6K (self-made, 1:1000), Histone H3 (Abcam #ab1791, 1:10000) and Histone H4 (Active motif #39269, 1:3000). HRP-conjugated secondary antibodies (Invitrogen) were used. Blots were developed using the ECL detection system (Amersham). Immunoblots were analysed using Image Lab program (Bio-Rad laboratories).

### RNA isolation and quantitative RT-PCR

Tissue was dissected, frozen on dry ice and stored at −80°C. Total RNA from guts of 10 flies was extracted using TRIzol (Invitrogen) according to the manufacturer’s instructions. mRNA was reverse transcribed using random hexamers and the SuperScript III First Strand system (Invitrogen). Quantitative PCR was performed using Power SYBR Green PCR (Applied Biosystems) on a QuantStudio 6 instrument (Applied Biosystems) by following the manufacturer’s instructions.

### Lysotracker and Cyto-ID staining, imaging and image analysis

Lysotracker dye accumulates in low pH vacuoles, including lysosomes and autolysomes, and Cyto-ID staining selectively labels autophagic vacuoles. Combination of both gives a better assessment of entire autophagic process ^29,42^. For the dual staining, complete guts were dissected in PBS, and stained with Cyto-ID (Enzo Life Sciences, 1:1000) for 30 min, then stained with Lysotracker Red DND-99 (Thermofisher, 1:2000) with Hoechst 33342 (1mg/ml, 1:1000) for 3 min. For the experiment only with Lysotracker staining, guts were stained with Lysotracker Red and Hoechst 33342 directly after dissection. Guts were mounted in Vectashield (Vector Laboratories, H-1000) immediately. Imaging was performed immediately using a Leica TCS SP8 confocal microscope with a 20x objective plus 5x digital zoom in. Three separate images were obtained from each gut. Settings were kept constant between the images. Images were analysed by Imaris 9 (Bitplane).

### Immunohistochemistry and imaging of the *Drosophila* intestine

The following antibodies were used for immunohistochemistry of fly guts; primary antibody: Phospho-Histone H3 (Ser10) (Cell Signalling #9701, 1:200). Secondary antibody: Alexa Flour 594 goat anti-rabbit (A11012, 1:1000). Guts were dissected in PBS and immediately fixed in 4% formaldehyde for 30 min, and subsequently washed in 0.1% Triton-X / PBS (PBST), blocked in 5% BSA / PBST, incubated in primary antibody overnight at 4 °C and in secondary antibody for 1 h at RT. Guts were mounted in Vectashield, scored and imaged as described above. For dysplasia measurement, the percentage intestinal length was blind-scored from luminal sections of the R2 region of intestines. For gut cell size measurement, nearest-neighbour internuclear distance in the R2□region was measured from raw image flies using the measure function in Fiji (Image J); 20□distances per gut, n ≥ 6 □ guts per condition.

### Library preparation and 16S sequencing / data analysis

Flies were washed in ethanol, then midguts were dissected in single PBS droplets, and 20 guts pooled per replicate. DNA extraction was performed using the DNeasy Blood&Tissue Kit (Qiagen) following the manufacturer’s instructions for gram-positive bacterial DNA, and using 0.1mm glass beads and a bead beater for 45s at 30Hz. Library preparation was performed following Illumina’s 16S Metagenomic Sequencing Library Preparation guide, with the following alterations: 100ng initial DNA amount, reactions for V1-V2 primer pair, amplicon clean-up with GeneRead Size Selection Kit following the DNA library protocol, and BstZ17I digest + gel extraction between PCR reactions for V1-V2 amplicons (for Wolbachia sequence removal). Pooled libraries were sequenced to 100,000 reads/sample on a HiSeq 250bp. Analysis was performed after quality control and paired-end joining for V1-V2 using the Qiime 1 pipeline and the greengenes database, at a depth of 20,000 reads/sample. Remaining *Wolbachia* sequences were removed bioinformatically before further analysis. For total quantification, qPCR with V3-V4 primers was performed with extension time of 1min. For validation, *Acetobacter pomorum* absolute amount was quantified by qPCR using bacteria-specific primers.

### Quantification and statistical analysis

Statistical analyses were performed in Prism (Graphpad) or R (version 3.5.5) except for Log-rank test using Excel (Microsoft). Sample sizes and statistical tests used are indicated in the figure legends, and Tukey post-hoc test was applied to multiple comparisons correction. Error bars are shown as standard error of the mean (SEM). For box-and-whiskers plots, Median, 25th and 75th percentiles, and Tukey whiskers are indicated. The criteria for significance are: NS (not significant) p>0.05; * p<0.05; ** p<0.01 and *** p<0.001.

## Notes

### Competing Interest Statement

The authors have declared no competing interest.

