## Supplementary_Figures for "Sexual identity of enterocytes regulates rapamycin-mediated intestinal homeostasis and lifespan extension"

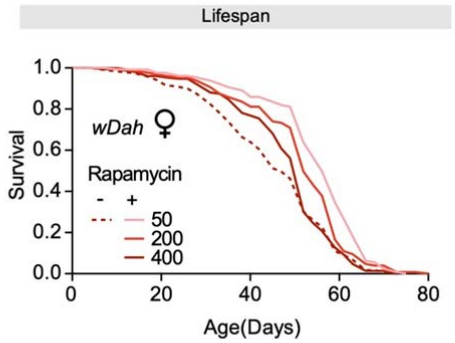


**Figure S1. Rapamycin treatment extends lifespan in *w*$^{Dah}$ females.**

Adult-onset rapamycin treatment in all three tested concentration (50, 200 and 400 µM) extended the median lifespan of *w*$^{Dah}$ females. (log-rank test, 50 µM p= 9.0E-08, 200 µM p=1.2E-03, 400 µM p=0.04, n >110 flies). See also Table S2.

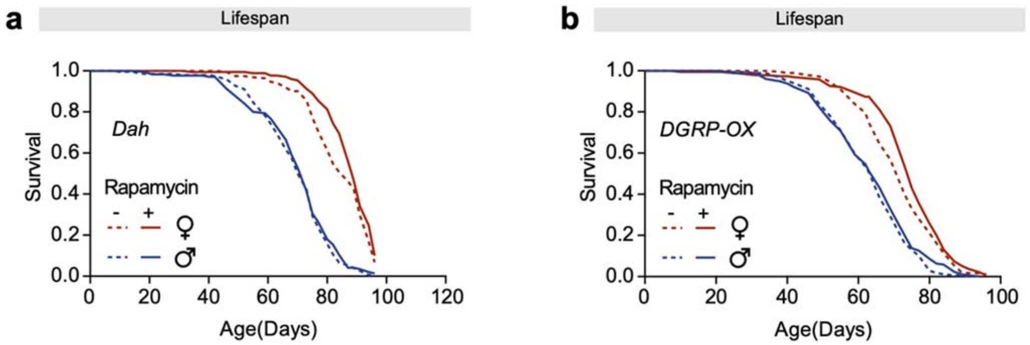


**Figure S2. Rapamycin treatment extends lifespan in *Dah* and *DGRP* females only.**

**a,** Adult-onset rapamycin treatment (200 μM) extended the lifespan of *Dah* females but not males (log-rank test, females $p=0.039$, males $p=0.73$, $n > 200$ flies). See also Table S3.

**b,** Adult-onset rapamycin treatment (200 μM) extended the lifespan of an outbred, genetically heterogenous fly line (*DGRP-OX*) females but not males (log-rank test, females $p=0.010$, males $p=0.23$, $n > 180$ flies). See also Table S3.

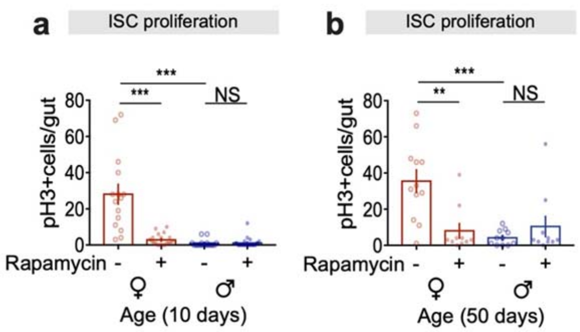


**Figure S3. Rapamycin treatment reduces ISC mitoses in females but not in males.**

**a,b,** Rapamycin treatment reduced the number of pH3 + cells in females, to the same degree as in untreated males, while it did not affect the number of pH3 + cells in males, (**a**) at 10 days of age, and (**b**) at 50 days of age (n ≥ 10 intestines, two-way ANOVA, interaction 10 days $p<0.001$, 50 days $p<0.01$; post-hoc test, NS $p>0.05$, **$p<0.01$, ***$p<0.001$).

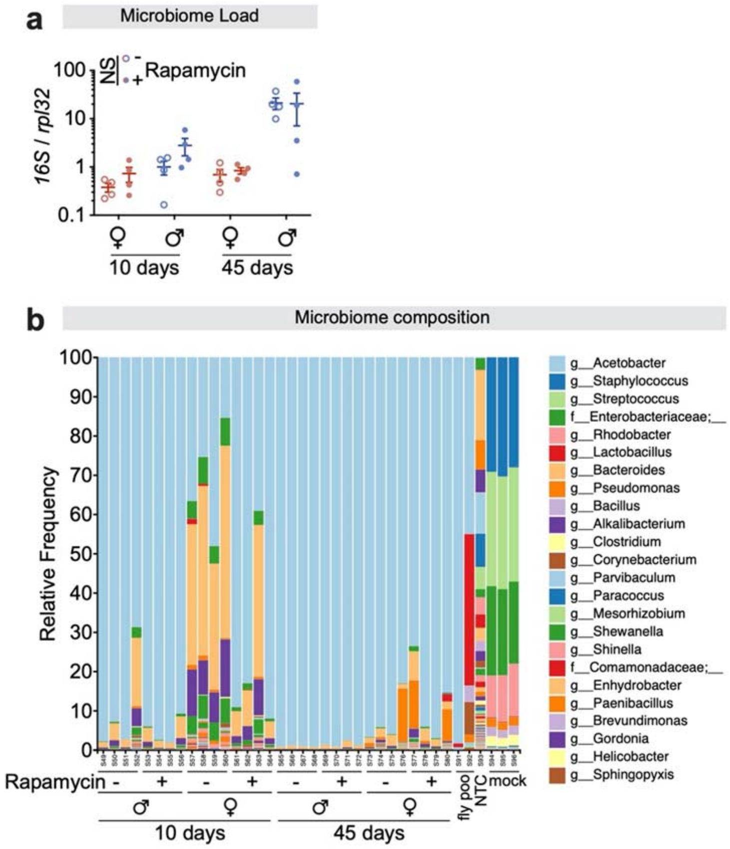


**Figure S4. The microbiome does not change upon treatment with rapamycin.**

**a,** Bacterial load in intestines changed with age and sex, but was not affected by rapamycin treatment (n = 4 biological replicates of 10 intestines per replicate, three-way ANOVA, age $p<0.001$, sexes $p<0.001$, treatment $p>0.05$).

**b,** Bacterial composition in intestines changed with age and sex, but was not affected by rapamycin treatment. (n = 4 biological replicates of 10 intestines per replicate, PERMANOVA (the number of permutations = 999), treatment $p>0.05$).

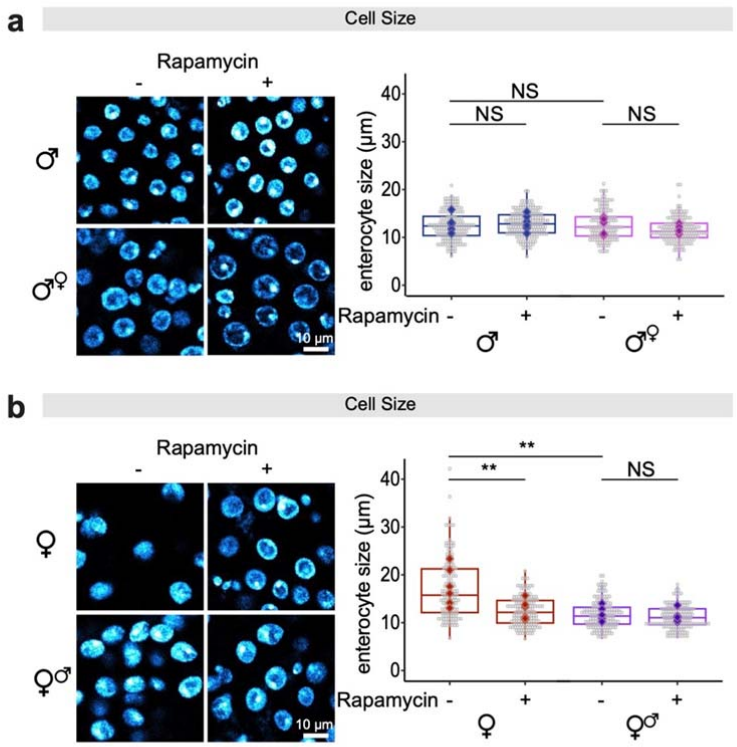


**Figure S5. Expression of *tra*$^{F}$ is necessary but not sufficient for the larger enterocyte size in female intestines.**

**a,** Expression of *tra*$^{F}$ in ECs in males did not affect cell size, neither did treatment with rapamycin (n = 6-8 intestines, n $\geq$ 20 enterocytes per intestine, circles indicate individual values and diamonds represent the average value per intestine; linear mixed model, interaction $p>0.05$; post-hoc test, NS $p>0.05$).

**b,** Knock-down of *tra*$^{F}$ in ECs in females reduced the size of enterocytes to the level of rapamycin-treated females, which was not further reduced by rapamycin treatment (n = 6-8 intestines, n $\geq$ 20 enterocytes per intestine, circles indicate individual values and diamonds represent the average value per intestine; linear mixed model, interaction $p<0.01$; post-hoc test, NS $p>0.05$, **$p<0.01$).

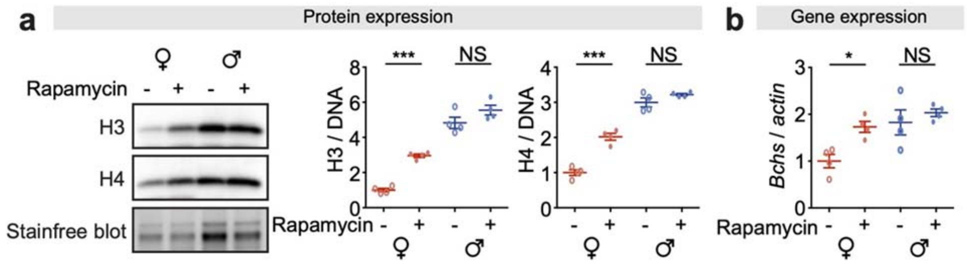


**Figure S6. Rapamycin treatment increased expression of histone H3, histone H4 and *Bchs* in intestines of females but not in males.**

**a,** The expression of histones H3 and H4 in intestines of females was lower than in males, and rapamycin treatment increased it (n = 4 biological replicates of 10 intestines per replicate, two-way ANOVA, H3, interaction $p<0.05$, H4, interaction $p<0.001$; post-hoc test, NS $p>0.05$, ***$p<0.001$).

**b,** The expression of *Bchs* in intestines of females was lower than in males, and rapamycin treatment increased it (n = 4 biological replicates of 10 intestines per replicate, two-way ANOVA, interaction $p>0.05$; post-hoc test, NS $p>0.05$, *$p<0.05$).
