## Supplementary_Tables for "Sexual identity of enterocytes regulates rapamycin-mediated intestinal homeostasis and lifespan extension"

**Table S1 - Rapamycin treatment extended lifespan in $w^{Dah}$ females but not in males (related to Figure 1)**

| Genotype/Treatment | Median Lifespan (days) | Maximum Lifespan (days) | n Dead | n Censored | % Increase (med) vs control (Control) | % Increase (max) vs control (Control) | Control | p-value (log rank) | | |
|---|---|---|---|---|---|---|---|---|---|---|
| | | | | | | | | female - Rapamycin | male - Control | male - Rapamycin |
| w ♀□ - Control | 65,5 | 88 | 138 | 5 | | | * | 2,07848E-06 | 1,70202E-24 | 1,21194E-21 |
| w ♀□ - Rapamycin | 70,1 | 88 | 152 | 1 | 7,0% | 0,0% | | | 3,21739E-37 | 2,0062E-34 |
| w ♂□ - Control | 51,6 | 78 | 167 | 4 | | | * | | | 0,77371499 |
| w ♂□ - Rapamycin | 51,6 | 78 | 153 | 1 | 0,0% | 0,0% | | | | |

**Cox Proportional Hazard (CPH) analysis**

| Details | Coefficient | Coefficient (estimate) | exp(coeff) | SE (coeff) | z | p |
|---|---|---|---|---|---|---|
| dead = 612 | Rapamycin | -0,531 | 0,588 | 0,123 | -4,32 | 1,60E-05 |
| censored = 11 | Sex | 1,172 | 3,230 | 0,120 | 9,78 | 2,00E-16 |
| | Rapamycin:Sex | 0,486 | 1,626 | 0,166 | 2,93 | 3,40E-03 |

**Table S2 - Rapamycin treatment extended lifespan in *w[Dah]* females but not in males (related to Figure 1 and S1)**

| Genotype/Treatment | Median Lifespan (days) | Maximum Lifespan (days) | n Dead | n Censored | % Increase (med) vs control (Control) | % Increase (max) vs control (Control) | Control | p-value (log rank) | | |
|---|---|---|---|---|---|---|---|---|---|---|
| | | | | | | | | w ♀ - Rapamycin 50 µM | w ♀ - Rapamycin 200 µM | w ♀ - Rapamycin 400 µM |
| w ♀ - Control | 46 | 70 | 118 | 0 | | | * | 9,00928E-08 | 0,001219441 | 0,039950423 |
| w ♀ - Rapamycin 50 µM | 55,1 | 75 | 126 | 1 | 19,8% | 7,1% | | | 0,019868832 | 8,77649E-08 |
| w ♀ - Rapamycin 200 µM | 50,4 | 70 | 127 | 2 | 9,6% | 0,0% | | | | 0,004560833 |
| w ♀ - Rapamycin 400 µM | 50,4 | 70 | 149 | 1 | 9,6% | 0,0% | | | | |
| | | | | | | | | w ♂ - Rapamycin 50 µM | w ♂ - Rapamycin 200 µM | w ♂ - Rapamycin 400 µM |
| w ♂ - Control | 43,5 | 68 | 130 | 1 | | | * | 0,60448407 | 0,75476739 | 0,995754772 |
| w ♂ - Rapamycin 50 µM | 41 | 68 | 116 | 2 | -5,7% | 0,0% | | | 0,445226822 | 0,722975386 |
| w ♂ - Rapamycin 200 µM | 43,5 | 77 | 122 | 1 | 0,0% | 13,2% | | | | 0,771149392 |
| w ♂ - Rapamycin 400 µM | 43,5 | 66 | 130 | 1 | 0,0% | -2,9% | | | | |

**Table S3 - Rapamycin treatment extended lifespan in *Dah* and *DGRP-OX* females but not in males (related to Figure S2)**

***Dah***

| Genotype/Treatment | Median Lifespan (days) | Maximum Lifespan (days) | n Dead | n Censored | % Increase (med) vs control (Control) | % Increase (max) vs control (Control) | Control | p-value (log rank) female - Rapamycin | male - Control | male - Rapamycin |
|---|---|---|---|---|---|---|---|---|---|---|
| Dah ♀ - Control | 85,5 | | 166 | 32 | | | * | 0,039377701 | 9,71687E-26 | 3,4958E-23 |
| Dah ♀ - Rapamycin | 88 | | 153 | 40 | 2,9% | | | | 2,52333E-38 | 5,21998E-35 |
| Dah ♂ - Control | 69 | | 168 | 41 | | | * | | | 0,725633352 |
| Dah ♂ - Rapamycin | 71,5 | | 148 | 40 | 3,6% | | | | | |

***DGRP-OX***

| Genotype/Treatment | Median Lifespan (days) | Maximum Lifespan (days) | n Dead | n Censored | % Increase (med) vs control (Control) | % Increase (max) vs control (Control) | Control | p-value (log rank) female - Rapamycin | male - Control | male - Rapamycin |
|---|---|---|---|---|---|---|---|---|---|---|
| DGRP ♀ - Control | 71 | | 182 | 7 | | | * | 0,016663578 | 2,14255E-10 | 8,54343E-06 |
| DGRP ♀ - Rapamycin | 74 | | 186 | 2 | 4,2% | | | | 1,07201E-18 | 1,62278E-11 |
| DGRP ♂ - Control | 64,5 | | 178 | 4 | | | * | | | 0,234867711 |
| DGRP ♂ - Rapamycin | 64,5 | | 174 | 7 | 0,0% | | | | | |

**Table S4 - Knock-down of Atg5 in ECs shortened the lifespan only in males (related to Figure 3)**

| Genotype/Treatment | Median Lifespan (days) | Maximum Lifespan (days) | n Dead | n Censored | % Increase (med) vs control (Control) | % Increase (max) vs control (Control) | Control | p-value (log rank) | | |
|---|---|---|---|---|---|---|---|---|---|---|
| | | | | | | | | 5966GS>Atg5 [RNAi] ♀□- RU486 | 5966GS>Atg5 [RNAi] ♂□- Control | 5966GS>Atg5 [RNAi] ♂□- RU486 |
| 5966GS>Atg5 [RNAi] ♀□- Control | 67 | 85 | 188 | 11 | | | * | 0,800188548 | 1,73274E-25 | 3,60157E-42 |
| 5966GS>Atg5 [RNAi] ♀□- RU486 | 67 | 81 | 187 | 12 | 0,0% | -4,7% | | | 9,27118E-25 | 2,19098E-40 |
| 5966GS>Atg5 [RNAi] ♂□- Control | 56 | 81 | 174 | 25 | | | * | | | 0,004539336 |
| 5966GS>Atg5 [RNAi] ♂□- RU486 | 52 | 78 | 192 | 7 | -7,1% | -3,7% | | | | |

**Cox Proportional Hazard (CPH) analysis**

| Details | Coefficient | Coefficient (estimate) | exp(coeff) | SE (coeff) | z | p |
|---|---|---|---|---|---|---|
| dead = 741 | sex | 1,1378 | 3,1199 | 0,1097 | 10,37 | 0,00000 |
| censored = 55 | RU486 | 0,00491 | 1,0503 | 0,1043 | 0,47 | 0,638 |
| | RU486:sex | 0,3004 | 1,3503 | 0,1481 | 2,03 | 0,043 |

**Table S5 - Overexpression of TraF in ECs restored the lifespan extention by rapamycin treatment in males (related to Figure 6)**

| Genotype/Treatment | Median Lifespan (days) | Maximum Lifespan (days) | n Dead | n Censored | % Increase (med) vs control (Control) | % Increase (max) vs control (Control) | Control | p-value (log rank) | | |
|---|---|---|---|---|---|---|---|---|---|---|
| | | | | | | | | mexG4>w - Rapamycin | mexG4>TraF - Control | mexG4>TraF - Rapamycin |
| mexG4>w - Control | 57 | 82 | 155 | 43 | | | * | 0,16950250337 | 0,854170991 | 2,25808E-05 |
| mexG4>w - Rapamycin | 57 | 79 | 181 | 18 | 0,0% | -3,7% | | | 0,092804711 | 0,000127203 |
| mexG4>TraF - Control | 57 | 79 | 141 | 58 | | | * | | | 1,55333E-06 |
| mexG4>TraF - Rapamycin | 63 | 82 | 114 | 85 | 10,5% | 3,8% | | | | |

**Cox Proportional Hazard (CPH) analysis**

| Details | Coefficient | Coefficient (estimate) | exp(coeff) | SE (coeff) | z | p |
|---|---|---|---|---|---|---|
| dead = 591 | Rapamycin | -0,135 | 0,8737 | 0,1104 | -1,22 | 0,2212 |
| censored = 204 | TraF | 0,0332 | 1,0337 | 0,1173 | 0,28 | 0,7774 |
| | Rapamycin:TraF | -0,4807 | 0,6184 | 0,1716 | -2,80 | 0,0051 |

**Table S6 - Kockdown of TraF in ECs blocked the lifespan extention by rapamycin treatment in females (related to Figure 6)**

| Genotype/Treatment | Median Lifespan (days) | Maximum Lifespan (days) | n Dead | n Censored | % Increase (med) vs control (Control) | % Increase (max) vs control (Control) | Control | p-value (log rank) | | |
|---|---|---|---|---|---|---|---|---|---|---|
| | | | | | | | | mexG4>v - Rapamycin | mexG4>TraF [RNAi] - Control | mexG4>TraF [RNAi] - Rapamycin |
| mexG4>v - Control | 71 | 86 | 196 | 3 | | | * | 1,81703E-09 | 1,56724E-09 | 7,72704E-12 |
| mexG4>v - Rapamycin | 76 | 91 | 184 | 15 | 7,0% | 5,8% | | | 0,915054013 | 0,466903337 |
| mexG4>TraF [RNAi] - Control | 76 | 89 | 197 | 2 | | | * | | | 0,381255525 |
| mexG4>TraF [RNAi] - Rapamycin | 76 | 89 | 190 | 9 | 0,0% | 0,0% | | | | |

**Cox Proportional Hazard (CPH) analysis**

| Details | Coefficient | Coefficient (estimate) | exp(coeff) | SE (coeff) | z | p |
|---|---|---|---|---|---|---|
| dead = 767 | Rapamycin | -0,652 | 0,521 | 0,104 | -6,25 | 4.2E-10 |
| censored = 29 | TraF [RNAi] | -0,626 | 0,535 | 0,102 | -6,12 | 9,50E-10 |
| | Rapamycin:TraF [RNAi | 0,553 | 1,739 | 0,145 | 3,81 | 0,00014 |
